# Real-world Interaction Networks Buffer Impact of Small Evolutionary Shifts On Biodiversity

**DOI:** 10.1101/013086

**Authors:** Gabriel E. Leventhal, Liyu Wang, Roger D. Kouyos

**Affiliations:** Institute of Integrative Biology, ETH Zurich, Switzerland; Institute of Robotics and Intelligent Systems, ETH Zurich, Switzerland; Department of Ecology and Evolutionary Biology, Princeton University, Princeton, NJ, USA; Division of Infectious Diseases and Hospital Epidemiology, University Hospital 9 Zurich, Zurich, Switzerland

**Author notes:** Corresponding author*: Gabriel Leventhal, Institute of Integrative Biology, Universitätstrasse 16, 14 8092 Zurich, Switzerland. These authors contributed equally to this work (Gabriel E. Leventhal, Liyu Wang).

**Keywords:** mutualistic networks, ecosystem stability, community diversity

## Abstract

Biodiversity maintenance and community evolution depend on the species interaction network. The “diversity-stability debate” has revealed that the complex interaction structure within real-world ecosystems determines how ecological communities respond to environmental changes, but can have opposite effects depending on the community type. Here we quantify the influence of shifts on community diversity and stability at both the species level and the community level. We use interaction networks from 19 real-world mutualistic communities and simulate shifts to antagonism. We demonstrate that both the placement of the shifting species in the community, as well as the structure of the interaction network as a whole contribute to stability and diversity maintenance under shifts. Our results suggest that the interaction structure of natural communities generally enhances community robustness against small ecological and evolutionary changes, but exacerbates the consequences of large changes.

## Author Summary

Ecological interaction networks are important determinants of the stability of a community and influence how the community responds to environmental changes. Network properties that stabilize antagonistic networks destabilize mutualistic networks and vice versa. Previous studies of interactions networks all assume that the type of interaction between species remains the same over time. Interactions between species, however, can shift from mutualistic to antagonistic and back over evolutionary time. Here we quantify the influence of such shifts on community diversity and stability at both the species level and the community level. We show that the location of the shifting species in the community, as well as the structure of the interaction network as a whole contribute to stability under such shifts. Our results suggest that the interaction structure of real-world communities generally enhances community robustness against small ecological and evolutionary changes, but exacerbates the consequences of large changes.

## 1 Introduction

Forty years ago Robert May theoretically predicted that large complex ecosystems with random interactions are unlikely to be stable [1]. Ever since then, a large body of work has been published investigating various aspects of community structures that can stabilise different types of ecological communities [2], [3]. Typically such studies have focused on two types of interactions. The first type is antagonistic interaction, such as in producer-consumer or predator-prey communities [4]–[6]. These interactions are characterised by one of the species benefiting from the interaction while the other species suffers a cost. The second type is mutualistic interaction, such as plant-pollinator or plant-seed disperser networks [7]–[10]. Here both interaction partners benefit from the interaction. Interestingly, the network properties that convey stability to antagonistic networks render mutualistic networks unstable and those that make mutualistic networks more stable make antagonistic networks less stable [11]. Recent studies on theoretical and real-world ecological networks containing both types of interactions showed that a low ratio of mutualistic to antagonistic interactions can destabilize an otherwise stable antagonistic community, but a moderate mixture of both interaction types can stabilize population dynamics [12]–[14].

These previous studies all assumed that the type of interaction between species remains the same over time. However, empirical studies have suggested that interaction types within an ecosystem shift over ecological and evolutionary time [15], [16]. It remains unclear what effect these interaction type shifts may have on the diversity and stability of an ecosystem. Here we perform a first step towards understanding how shifts between interaction types in real-world communities affect biodiversity and community stability at both the species level and the community level. Mutualists are continuously under the threat of exploitation [17], and therefore shifts from mutualism to antagonism in such mutualistic communities represent a relevant case study for the effect of shifts in ecological communities [16].

More specifically, we examined 19 previously published real-world plant-pollinator networks [7] and applied a species interaction model to study the change in the equilibrium state of the community. The general strategy was to choose one or several pollinator species in each network and switch their interactions with all plants to antagonistic. We determine the effects of these shifts with respect to resistance stability [18], evenness biodiversity [19] and ecological dominance on the shifted species. As proxies for these properties we use the relative Eu-clidian distance, the relative Shannon Index change, and the relative frequency change of the shifted species between the two equilibrium states respectively (see Methods). To assess the importance of network structure, we compared each real-world network with 100 simulated networks generated by randomizing the interactions.

## 2 Materials and Methods

We determine the effects of shifts of the interaction type in simulated real-world networks (see Section 2.1). A network is composed of a number of pollinator and plant species. A specific pollinator only interacts with a subset of plant species (see Figure 1a). This network of interactions is represented by an adjacency matrix. The strength of the interaction is given by the interaction model (see Section 2.2).

We study the stability of the real-world networks on four different levels. In a first step, we examine the effect of the shift of the interaction type of a single species on the network as a whole, as well as on the shifting species itself (see Section 2.4). This allows us to quantify the effect of the shifting species’ centrality in the network (see Section 2.5) on diversity and stability. We also examine the effect of the species centrality on the change of its own abundance. In a second step, we compare the effect of shifting random species in a real-world network to equivalent randomized networks using two randomization schemes (see Section 2.6). The first randomization scheme keeps the connectance constant, while destroying all network structure. The second scheme keeps both the connectance and the degree distribution constant. This allows us to tease apart the effects of connectance, degree distribution and higher order network structure. To quantify the effects of the introduction of an antagonistic interaction on the network, in a third step, we study the differences between the interaction type shift of a single species and the extinction of the same species. Finally, in a fourth step, we determine the robustness of the network to interaction type shifts of multiple species. As for single species, we quantify the stability and diversity of the network to such shifts. For multiple shifts, we also quantify the number of secondary extinctions caused by the interaction type shifts.

In general, all simulations proceed in the following manner. We first determine the equilibrium abundances of each species in the network as reference abundances of a specific network(see Section 2.3). We then shift the interaction type of a single pollinator species. This results in a new interaction network (see Figure 1b). Finally, we determine the new equilibrium abundances of each of the species and compare these to the reference abundances.

**Figure 1:**
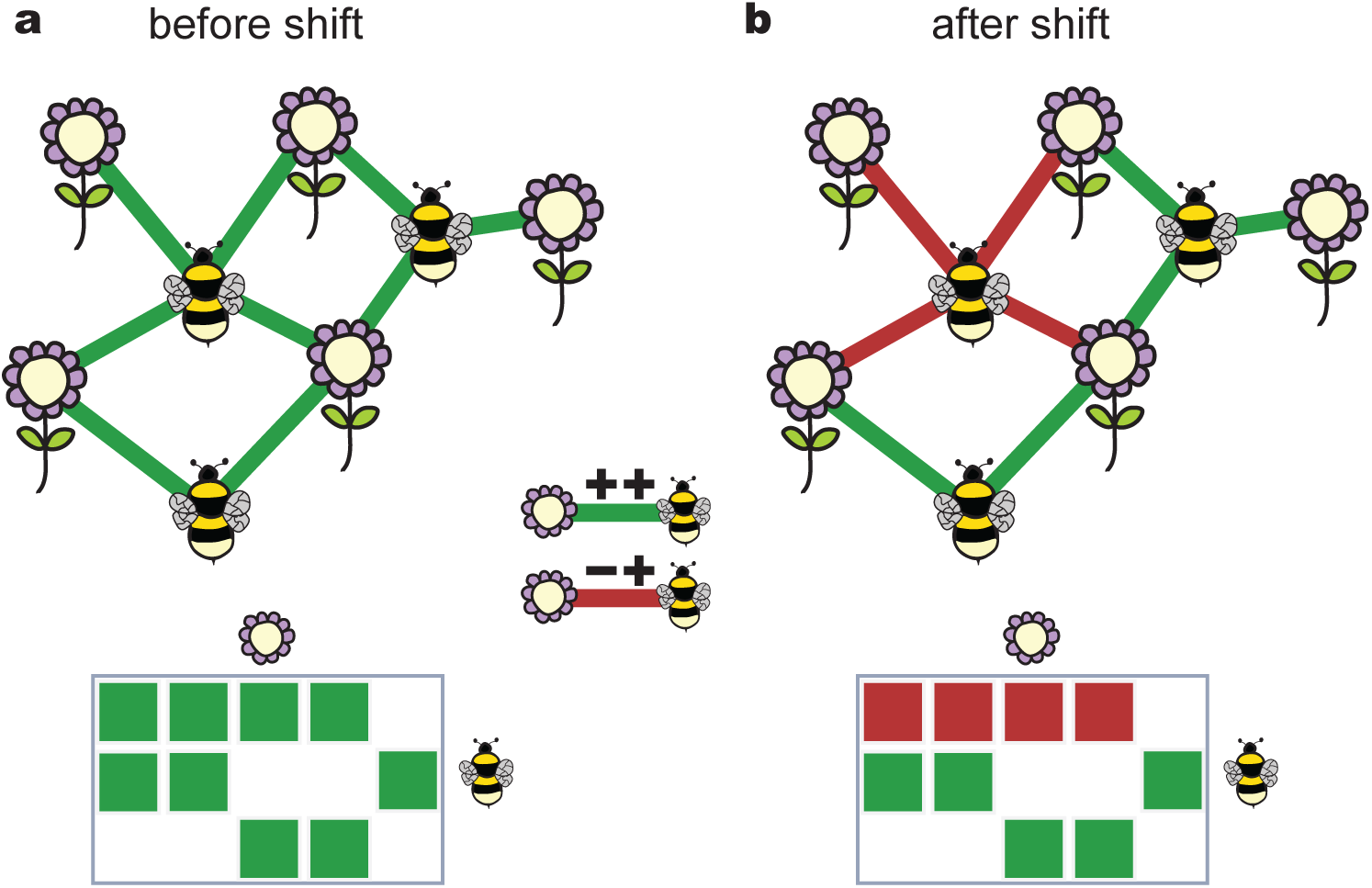
Schematic of a mutualistic network that undergoes an interaction type shift. Before the shift, all interactions between plants and pollinators are beneficial for both the plant and the pollinator (green: ++ mutualistic interaction). After a pollinator shifts, interaction with plants remain beneficial to the pollinator, but are detrimental to the plant (red: −+ antagonistic interaction). This is modeled by inverting the signs of the interactions between the pollinator and the plants in the adjacency matrix of the network.

### 2.1 Network data

We use real world plant-pollinator networks previously published in [7]. The smallest network has 10 plants and 12 pollinators, while the largest has 112 plants and 839 pollinators. A full list of number of species in each network is given in Supplementary Table S1. The values of interaction strength between plants and pollinators have been normalized such that bidirectional asymmetry exists [7].

**Table 1:** Rank correlation (Spearman’s *ρ*) between the centrality measures and the distance measures in the case of single shifts within each network. Parameters: *r*_*i*_ = 0:4; *s*_*i*_ = 1:5; *c*_*i*_ = 1. A double asterisk signifies *p* < 0:01, a single asterisk *p* < 0:05.

| Network | Betweenness centrality |  |  | Degree centrality |  |  |
| --- | --- | --- | --- | --- | --- | --- |
| | $\delta$ | $\vert \Delta s\vert $ | $\Delta f$ | $\delta$ | $\vert \Delta s\vert $ | $\Delta f$ |
| 1 | 0.861** | 0.819** | −0.626** | 0.881** | 0.845** | −0.652** |
| 2 | 0.709** | 0.658** | −0.416** | 0.707** | 0.658** | −0.41** |
| 3 | 0.617** | 0.597** | −0.445** | 0.597** | 0.578** | −0.422** |
| 4 | 0.63** | 0.609** | −0.434** | 0.623** | 0.6** | −0.417** |
| 5 | 0.714** | 0.692** | −0.535** | 0.737** | 0.714** | −0.557** |
| 6 | 0.63** | 0.617** | −0.427** | 0.632** | 0.62** | −0.428** |
| 7 | 0.679** | 0.656** | −0.427** | 0.679** | 0.657** | −0.423** |
| 8 | 0.612** | 0.6** | −0.438** | 0.606** | 0.594** | −0.426** |
| 9 | 0.684** | 0.666** | −0.465** | 0.693** | 0.674** | −0.47** |
| 10 | 0.563* | 0.564* | −0.219 | 0.567* | 0.567* | −0.212 |
| 11 | 0.592** | 0.55** | −0.368* | 0.602** | 0.564** | −0.374* |
| 12 | 0.843** | 0.755** | −0.749* | 0.867** | 0.791** | −0.788** |
| 13 | 0.705** | 0.684** | −0.395** | 0.722** | 0.698** | −0.403** |
| 14 | 0.955** | 0.944** | −0.908** | 0.963** | 0.955** | −0.912** |
| 15 | 0.741** | 0.688** | −0.509** | 0.737** | 0.686** | −0.514** |
| 16 | 0.679** | 0.646** | −0.484** | 0.677** | 0.645** | −0.472** |
| 17 | 0.747** | 0.743** | −0.611** | 0.751** | 0.746** | −0.607** |
| 18 | 0.83** | 0.786** | −0.747** | 0.834** | 0.789** | −0.749** |
| 19 | 0.598** | 0.593** | −0.415** | 0.595** | 0.59** | −0.407** |

### 2.2 Interaction Model

We use a multi-species interaction model equivalent to Bascompte *et al*. [7]. For mathematical tractability, we focus on Holling type I functional responses and show in the Supplementary Materials that we obtain qualitatively equivalent results in the case of single species shifts when using a Holling type II functional response. The model with a type I functional response in the interaction term has the form,

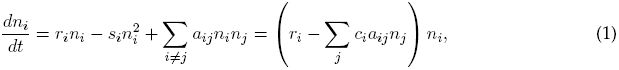

where *n*_*i*_ is the abundance of species *i*, *r*_*i*_is species’ *i*’s individual growth rate, *s*_*i*_ a intra-specific competition term and *a*_*ij*_ is the effect on species i of the interaction with species *j*. We have incorporated the intra-specific interaction term for a species i into the interaction matrix, such that *a*_*ii*_ = *s*_*i*_. The parameter *c*_*i*_ represents the magnitude of the payoff of an interaction. Positive entries in A (*a*_*ij*_ > 0) are interactions where the species *i* pays a price for interacting with *j* and negative entries (*a*_*ij*_ < 0) are interactions where species *i* gains a benefit from the interaction with species *j*. Note that the interaction matrix A contains both the effects on plants when they interact with a animals, as well as the effects on animals, when they interact with plants.

In this study, we only consider interactions between two species classes (plants and pollinators). Assuming there are *N* species in total, of which *N*_*A*_ are animals species (pollinators) and *N*_*P*_ are plant species, then we can partition the matrix A such that the rows and columns *i* ∈ {1, 2, …, *N*_*A*_} = *N*_*A*_ are animal species and *i* ∈ {*N*_*A*_ + 1, *N*_*A*_ + 2, …, *N*_*A*_} = *N*_*P*_ are plant species. We neglect inter-specific competition (plant-plant and animal-animal interactions), so *a*_*ij*_ = 0 when *i; j* ∈ *N*_*X*_, *X* ∈ {*N*_*A*_, *N*_*P*_}.

To shift the *k*-th animal from a mutualist into an antagonist, we multiply the *k*-th column of A by −1. Therefore, the animal still gains the same benefit from interacting with the plants as before the shift, but the plant now pays a price for each interaction with this animal species. In terms of a plant-pollinator system, if the interaction strengths are related to the frequency of visits, then the pollinator still claims the reward for visiting the plant (e.g. gets nectar), but the plant pays a price for each visit (e.g. damaging of the plant, theft of nectar).

We assume that the frequency of visitation is equally proportional to the abundance of any pollinator species. Thus generalist pollinators distribute their mutualistic interactions across many different plants, while specialist pollinators visit few plants more frequently. Mathematically, we assume that the columns of A excluding the diagonal sum to 1,

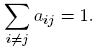

We consider facultative mutualism, such that *r*_*i*_ > 0 and all species have a positive growth rate in absence of any mutualistic interaction. Population growth is limited by the intra-specific competition terms *s*_*i*_. For randomly interacting populations, a finite non-trivial steady state for species i exists if *s*_*i*_ > *c*_*i*_ (see Supplementary Materials). We show in the Supplementary Materials that we obtain qualitatively equivalent results with obligate mutualism.

### 2.3 Determining the equilibrium abundances

We find the non-trivial equilibrium state of a system with interaction matrix A by solving the nonlinear system of equations *d*n/*dt* = 0 (Equation 1). This is equivalent to finding the solution to the linear system of equations 
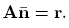
 = r.

A species *i* is extinct at equilibrium if it has an equilibrium abundance 
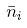
 ≤ 0. These negative abundances, however, can influence the equilibrium abundances of the other species. We therefore extract a sub-matrix A— where the *i*-th row and column are deleted and find the equilibrium abundances of this subsystem n′*. We repeat this procedure until all remaining species have an abundance 
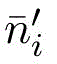
 > 0.

### 2.4 Stability and diversity measures

We use three measures to determine the effect of interaction shifts on the stability and diversity of the network: (1) relative Euclidian distance; (2) relative frequency change of the shifting animal; and (3) the relative Shannon index change of the equilibria.

#### Relative Euclidian distance

The relative Euclidian distance *δ* measures how much the abundances of the different species change from before to after the shift of one or more animal species from mutualists to antagonists. If *n*_*i*_ is the abundance at equilibrium of species *i* before the shift and 
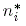
 is the abundance at equilibrium of species i after the shift, then the relative Euclidian distance is,

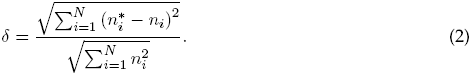

#### Relative frequency change of the shifting animal

The relative change in an animal’s frequency ∆*f*_*i*_ measures the relative benefit or price an animal gains or pays if that animal shifts from being a mutualist to an antagonist,

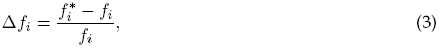

where *f*_*i*_ is the relative of abundance of the animal species in the current ecosystem,

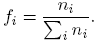

#### Relative Shannon index change

The relative Shannon index change **∆***s* measures how much diversity is affected by the shift of one or more animals from pure mutualists to antagonists,

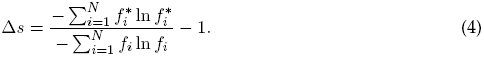

### 2.5 Network measures

Centrality measures determine the importance of an individual species in the network. Here we use two centrality measures that capture different aspect of the species placement in the network [20].

#### Degree centrality

We measure the importance of single species within one network by its degree centrality *k*_*i*_, which is defined as the number of interactions *K*_*i*_ with other species divided by the total number of species,

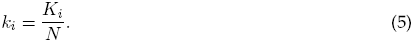

#### Betweenness centrality

An alternative measure of centrality of a single species is its betweenness. The betweenness of species *i* is defined as the number of shortest paths between any two nodes *l* and *m* that pass through *i* divided by the total number of shortest paths between *l* and *m*. We calculate betweenness centrality using the algorithm implemented in the igraph library [21].

#### Nestedness measure

We measure nestedness using the established NODF measure [22]. In order to account for network properties such as the total number of interaction that can also influence nestedness, we use a relative measure of nestedness, 

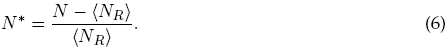

Here 
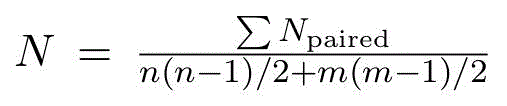
 is the NODF measure defined in [22] and <*N*_*R*_> is the NODF nestedness of an interaction matrix obtained by shuffling the interactions, averaged over 100 randomizations (see section 2.6).

### 2.6 Randomizations

We compare the effects of shifting from mutualists to antagonists in the real network to randomized networks with equal number of plant and animal species. The randomized networks have the same number of plant-animal interactions. We employ two randomization schemes. In the first scheme, all interactions are distributed randomly between plants and animals. To guarantee that all animals interact with at least one plant and vice-versa, a single interaction partner is first assigned to each animal/plant. The remaining interactions are then randomly placed to animal-plant pairs. This randomizes both the degree distribution as well any other higher-order structure of the network. In the second scheme, the degrees of each animal and plant are retained, but the interaction partners are randomized. In this scheme, only the higher-order structure is randomized.

### 2.7 Residual measures.

When species are removed from the community (i.e. extinctions) their abundances are artificially set to zero. This is a different perturbation than shifts to antagonism, since in shifts the interactions are modified and in extinctions the abundances are modified. In such cases, the distance measures may be artificially inflated due to the removal of a single species. For example, if species *k* is removed it’s abundance is set to zero, while if it shifts it may persist with a non-zero abundance, leading to a smaller change in the case of shifts. We therefore consider residual measures of δ, ∆*s* and ∆*f*, which are calculated equivalently to explained above without considering the shifted/removed species,

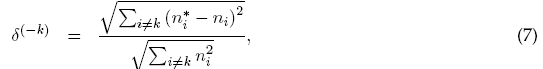

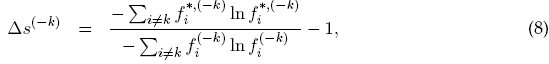

where

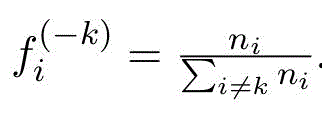

## 3 Results

### 3.1 Single shifts

#### Response to single shifts

Shifting the interaction type of a single pollinator species changed the equilibrium abundances of all the species but did not result in any secondary extinctions of either plants nor pollinators. As a result of the introduction of antagonistic interactions, however, the total sum over the abundance of all species decreases. The relative Euclidian distance, *δ*, between the equilibrium states before and after the single shift is smallest when the shifting species has a low centrality and increases with the centrality of the species (Figure 2 and Table 1). The relative Shannon index change, ∆*s*, is always negative for single shifts. The magnitude of the relative Shannon index change, |∆*s*|, is smallest for low centrality species and increases with species centrality. The relative abundance frequency change of the shifting species is either positive or negative. It is largest for low centrality species and decreases with increasing centrality. The rank correlation between the centrality and both *δ* and |∆*s*| is significantly positive within each network and independent of the centrality measure (Table 1). We also qualitatively find the same correlations between the centrality and distance measures for weak and strong intraspecific competition (see Supplementary Materials).

**Figure 2:**
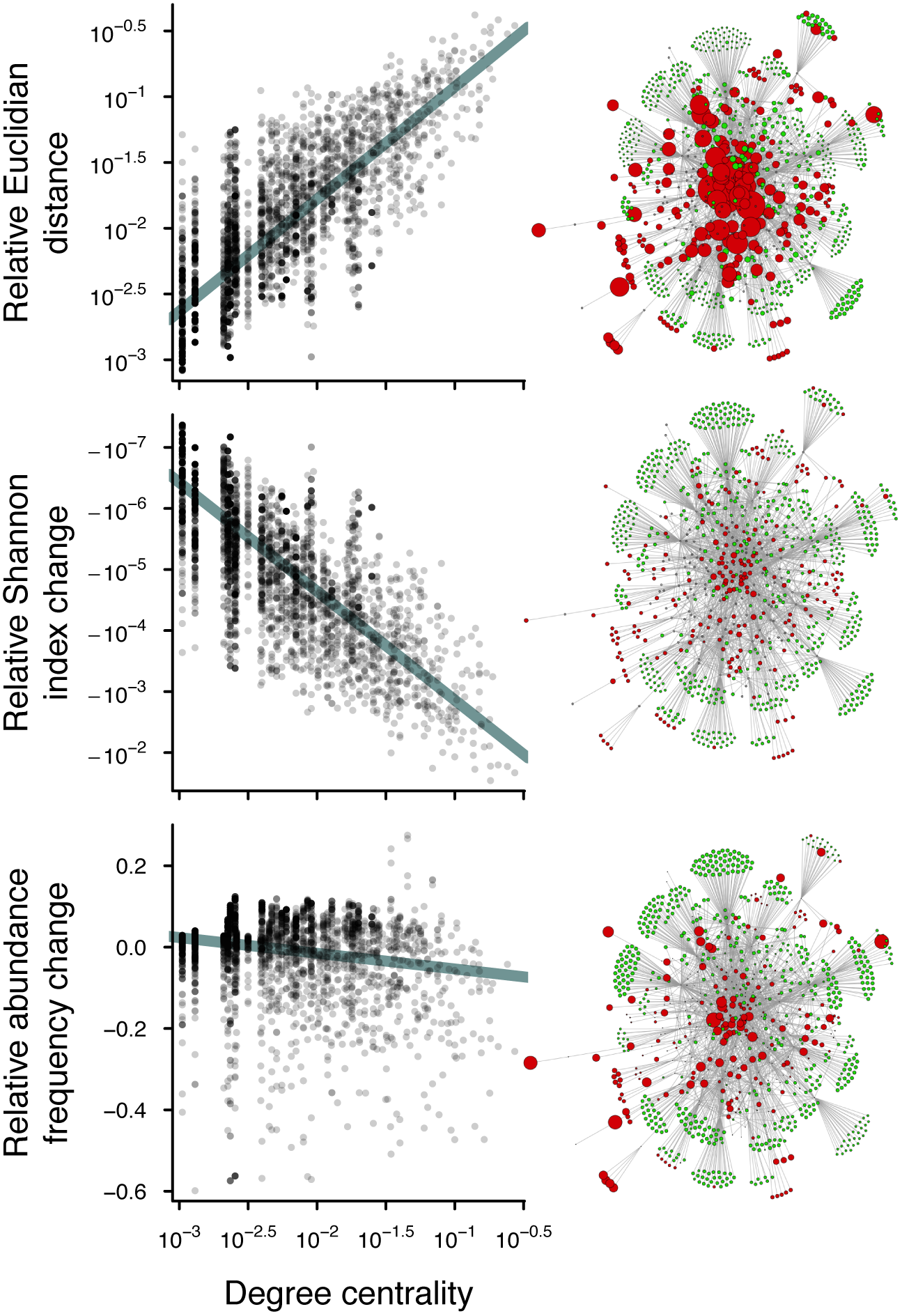
Effect of degree centrality on the different stability measures. Relative Euclidian distance change (top), relative Shannon index change (middle) and relative frequency change (bottom) as a function of the degree centrality of the shifting species for all networks combined. Each point represents a single shifted pollinator species. The model parameters are: *s*_*i*_ = 1:5; *r*_*i*_ = 0:4; *c*_*i*_ = 1. The networks figures to the right of the plots give a qualitative impression of the on the dependence of the measures on centrality for the plant-pollinator dataset from Kibune Forest, Kyoto, Japan [7], [32]. Grey nodes correspond to plant species and coloured nodes to pollinator species. Green animal species increase their relative abundance when shifting to antagonism and red species decrease their relative abundance. The size of the nodes is proportional to the magnitude of the respective measure.

#### Real networks versus randomizations

To asses the effect of network structure of the stability to shifts, we compared the median value of the three distance measures for all possible single species shifts in real-world and corresponding randomized networks (mean value over 100 randomizations). The median relative Euclidian distance and the median magnitude of the relative Shannon index change are higher in the randomized networks than the real-world networks (Figure 3a-b). Conversely, the relative frequency change of the antagonistic species is generally larger in real-world than in randomized networks (Figure 3c). These results are consistent for both randomization schemes, though the effect is smaller when the degree distribution is kept constant and only the higher order structure is randomized. Weak and strong interactions did not qualitatively influence these results (see Supplementary Materials).

**Figure 3:**
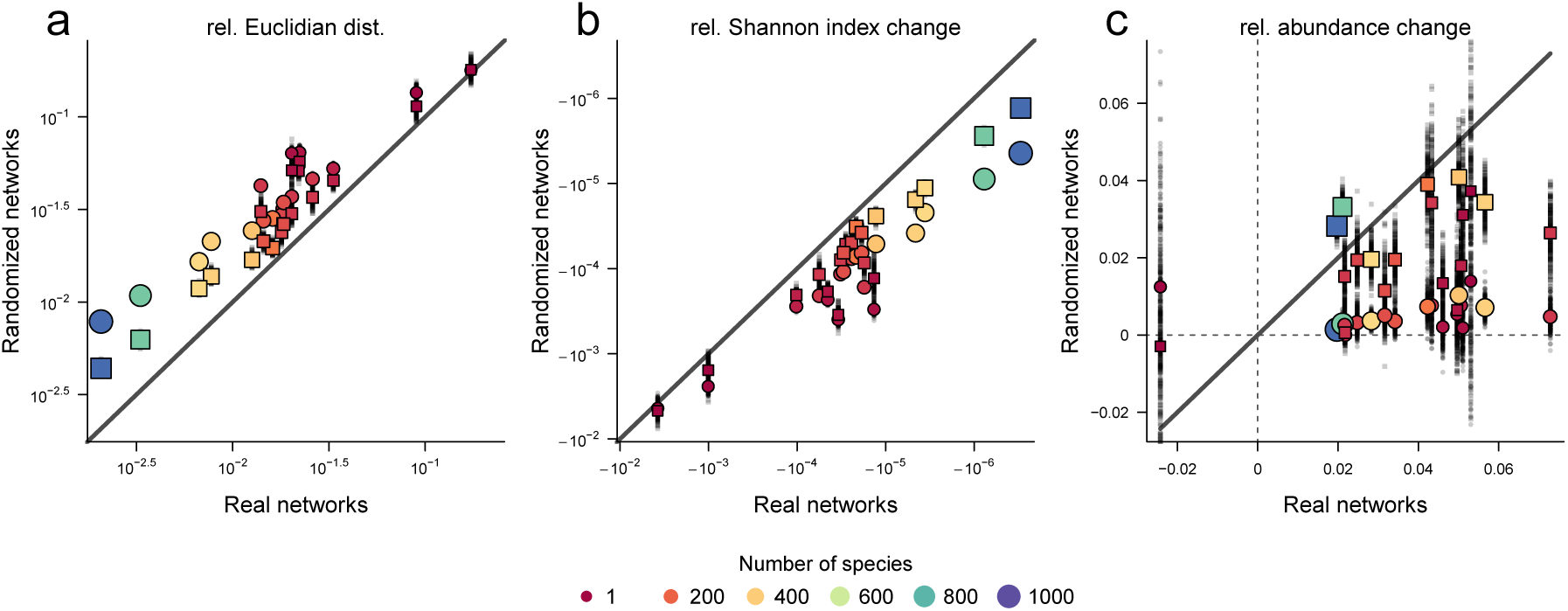
The mean values of the stability measures in the real networks vs. their randomized counterparts. The coloured circles and squares are the median values over all pollinators for each of the three measures and each network under the randomization scheme that discards and retains the degree distribution respectively. The position on the y-axis is the mean value of this median over 200 repetitions. The individual median values are indicated by small black dots (partially hidden behind the coloured circles). The size and colour of the circles are representative of the number of species in each network. Points above (below) the diagonal for positive (negative) values indicate larger values for randomized networks. The model parameters are the same as in Figure 2: *s*_*i*_ = 1:5; *r*_*i*_ = 0:4; *c*_*i*_ = 1. The effect of weak and strong intra-specific competition is shown in Supplementary Figures S5 and S6.

#### Influence of nestedness

We then determined the role of nestedness on the stability to shifts. In agreement with previous findings [8], networks with more species are more nested than networks containing less species (Supplementary Figure S1). We find that the median relative Euclidian distance decreases both with number of species (Spearman’s *ρ* = −0:984, *p* = 3:20 · 10^−14^) and nestedness (Spearman’s *ρ* = −0:753, *p* = 0:000305), but increases with connectance (Spearman’s *ρ*= 0:921, *p* = 1:50 · 10^−6^). The magnitude of the median relative Shannon index changes ∆*s* also decreases with number of species (Spearman’s *ρ* = −0:947, p = 7:86 · 10 ^−13^) and nestedness (Spearman’s ρ = −0:707, p = 0:00101), and also increases with connectance (Spearman’s *ρ* = 0:863, *p* < 10^−15^). The median relative frequency change ∆*f* increases with number of species (Spearman’s *ρ* = 0:548, *p* = 0:0152) and nestedness (Spearman’s *ρ* = 0:496, *p* = 0:0323), but decreases with connectance (Spearman’s *ρ* = −0:574, *p* = 0:0116).

Since, however, number of species, connectance and nestedness are not necessarily independent (Figure S1), we also perform an ANOVA with all three factors. When analyzed together, only the number of species (*F* = 75:8; *p* = 3 · 10^−7^, *df* = 1) and the connectance (*F* = 5:98; *p* = 0:0273, *df* = 1) are significant. The normalized NODF no longer has a significant effect (*F* = 0:147; *p* = 0:707, *df* = 1).

### 3.2 Shifts to antagonism versus extinctions

We compared the relative Euclidian distance and the relative Shannon index change when a single species shifts to antagonism to the case where the same species is removed, i.e. goes extinct (Figure 4). As the extinction of a species in itself represents a strong change in the distribution of species abundances, even without considering the secondary effects of this extinction on other species, we only consider the change in the species excluding the antagonistic/extinct species (Supplementary Matrials). Interaction type shifts affect the community more strongly than the removal of a mutualist, both with respect to relative Euclidian distance (*ρ* = −0:571; *p* < 10 ^−16^) and relative Shannon index change (*ρ* = −0:406; p < 10 ^−16^). Moreover, we find that the difference in effects between interaction type shifts and extinctions decreases with the centrality of the shifted/removed species. The effect of weak and strong intra-specific competition is shown in Supplementary Figures S7.

**Figure 4:**
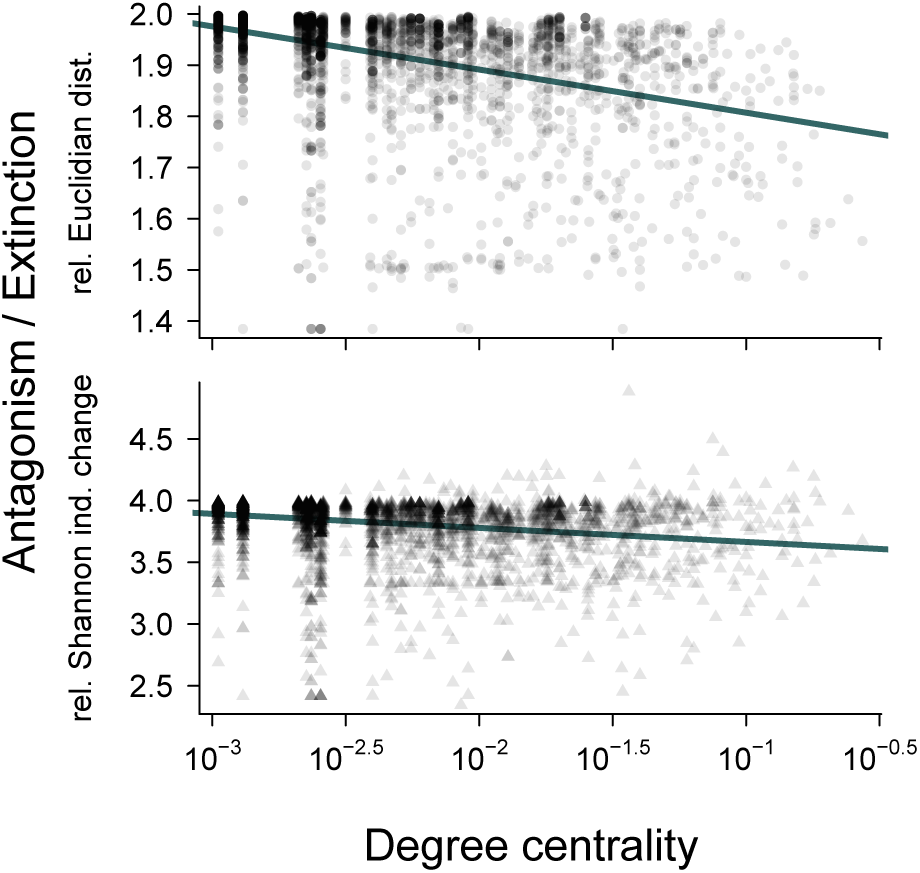
The effects of shifts to antagonism vs. extinctions of pollinators. Top panel: The relative Euclidian distance and relative Shannon index change when a pollinator shifts to antagonism divided by the measure when that same pollinator is removed, respectively. *s*_*i*_ = 1:5; *r*_*i*_ = 0:4; *c*_*i*_ = 1.

### 3.3 Multiple shifts

We then assessed how networks respond to shifts of multiple species. For a small number of shifts, both the relative Euclidian distance and the Shannon index change is smaller in real-world than randomized networks (similar to single species shifts). To this end we compared *δ* and ∆*s* in real and randomized networks when shifting different fraction of randomly chosen pollinators. As the number of shifted species increases, the effect on real-world networks becomes considerably larger than in randomized networks (Figure 5; we also observe this reversal in the case of species removals, see Supplementary Figure S3). When a sufficient number of pollinators are shifted, we started to observe secondary extinctions. The number of such secondary extinctions is considerably larger in the real-world networks than randomized networks (Supplementary Figure S2). We observe an initial increase in the number of secondary extinctions with the number of shifted species. However, when almost all species shift to antagonism then the number of secondary extinctions decreases again (Supplementary Figure S2).

**Figure 5:**
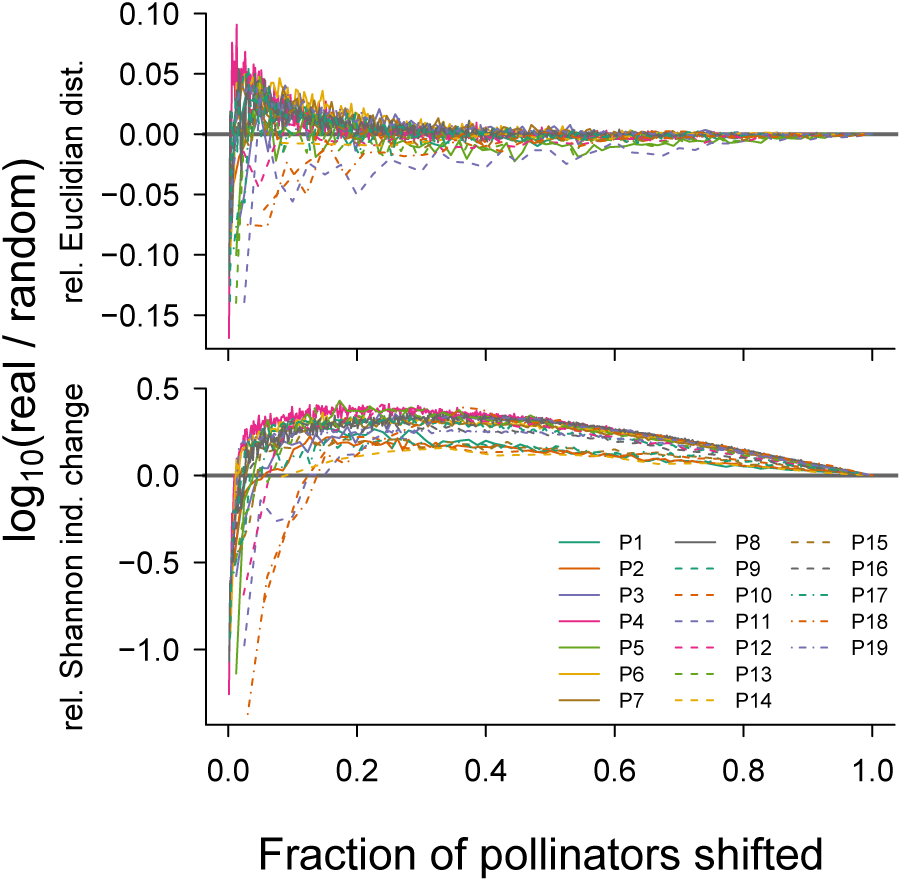
Multiple shifts to antagonism. The relative Euclidian distance and the relative Shannon index change as a function of the fraction of pollinators that shift to antagonism in real compared to randomized networks. The effect of only a small number of pollinator shifts is larger in randomized than real-world networks (negative values). As the number of shifting pollinators increases, this is generally reversed and the effects on the community become larger in real-world than randomized networks (positive values). For each network, we randomly sample *m* = 1; 2,…, *N*_*A*_ pollinators and shift their interaction to antagonism (*N*_*A*_ = number of pollinators). The plotted values are the mean over 200 such samples at each *m* for the both real-world network and 100 randomizations. *s*_*i*_ = 1:5; *r*_*i*_ = 0:4; *c*_*i*_ = 1.

## 4 Discussion

More centrally located species, as measured by a higher degree or betweenness centrality, pay a higher cost when shifting to antagonism, while species towards the periphery of the network, lower degree centrality, pay a smaller cost or even benefit from shifting (Figure 2). Similarly, the relative change in Shannon Index and the relative Euclidian distance increase with the centrality of the shifted species (Figure 2). This confirms the importance of generalists in maintaining diversity and stability [23], [24]. Together, this implies that shifts of species at the edge of the network are more likely to occur, both because of the advantage to the species itself, as well as the smaller effect on the community.

The effect of a single pollinator species shift to antagonism, however, also strongly depends on network structure (Figure 3). The total change in equilibrium abundances and species evenness is larger in randomized than in real-world networks. The relative effect on the shifting species, however, is mostly positive (i.e. the shifting species increases its abundance relative to all other species) and its magnitude is generally larger in real-world than in randomized networks. Overall, these results have three implications. Firstly, real-world mutualistic networks have an intrinsic structure that reduces the effect on the whole community caused by shifts to antagonism. Secondly, real-world networks, in contrast to randomized networks, maintain evenness biodiversity better under shifts. Thirdly, real-world networks enhance the benefit the shifting species receives compared to all the other species in the network.

These results are generally consistent when all network structure is destroyed, and also when the degree distribution of the network is kept constant, although the effect is smaller in the latter case. Thus, the degree distribution can only in part explain the difference between real-world and randomized networks. This is consistent with previous reports that the degree distribution can be a predictor of network stability, but its effect must be teased apart from the effects from other higher-order network structure [25], [26].

Mutualistic communities are often strongly nested [27], which could explain the difference between real-world and randomized networks that is not due to differences in the degree distribution. We found that the median values of the relative Euclidian distance and the magnitude of the relative Shannon index change both decrease with relative nestedness (Supplementary Figure S1). The effect of relative nestedness, however, becomes non-significant once community size is taken into account. Therefore it is unclear whether the robustness of real-world networks is a direct consequence of their nested architecture or other higher-order network structure.

An alternative perturbation to which the stability of an ecological community can be measured is the extinction or removal of a species [28], [29]. Interaction type shifts have a greater effect on the community than extintions. Even for the most centrally located species, interaction type shifts lead to more than 50% larger changes compared to extinctions. Thus although interaction type shifts lead to greater changes than removals, ecological communities are comparatively more robust to shifts in central species than to extinctions.

Even though the structure of real-world mutualistic networks enhances their robustness against single shifts, the opposite is the case for multiple shifts, i.e. they are more sensitive to multiple shifts than randomized networks. Mougi & Kondoh [14] previously showed that adding mutualistic interactions to predator-prey networks can stabilize the community, but that too many mutualistic interactions decreases the stability again. Here, we replace mutualistic interactions with anatagonistic ones and find that changing only a small number of interactions drastically destabilizes real-world networks. As more and more interactions are shifted, stability is regained. Equivalently, Allesina & Tang [13] showed that a large number of weak mutualistic interactions has a destabilizing effect on antagonistic communities. Thus, these results are in agreement, and the effect of mixed-effects is similar, whether one starts with naturally stable antagonistic networks or a naturally stable mutualistic network.

Due to the general nested architecture of real-world mutualistic communities [27], there are more specialists than generalists in these communities. These specialists have a lower degree centrality and are more likely to benefit from shifting their interaction type but will have less of an effect on the whole community. Thus the species that contribute most to the stability of the community (i.e. generalists) are also those which are most prohibited from shifting their interaction type. Generalist species have also been shown to be less tolerant to a decrease in the strength of the mutualistic interactions [30]. These results offer an additional explanation why in phylogenetic studies of mutualistic communities specialists are more likely to “disappear” from the community [16], [24]. While extinctions have been the usual explanation, we propose that interaction type shifts could also be a driver of this observation.

Exploitation represents an important challenge for mutualistic communities since cooperation between unrelated individuals is particularly susceptible to cheating [17], [31]. The effect of such cheating has been well studied in the case of two-species interactions, but it is not clear how these changes propagate through the complex interaction networks characterising real ecological communities. The interaction type shifts studied here can be seen as the result of successfully invading cheaters. Overall, our results indicate that real-world networks promote the shifting of a typical single species more strongly compared to random community assemblies, but these networks are structured in a way that buffers the effect of interaction type shifts of single species on the community as a whole. This increased resistance of real-world networks to single species shifts compared to random assemblies wanes and is eventually reversed as the number of shifting species increases. This indicates that the structure of real-world networks may protect these communities from small perturbations such as single species shifts, but can exacerbate the consequences of large perturbations such as multiple species shifts. This property of real-world networks might be especially relevant as the currently on-going anthropogenic changes may lead to the type of large-scale perturbations to which ecological communities are particularly sensitive according to our findings.

## Supplementary Figures

S1 The effect of number of species, connectance and relative nestedness on the median values of the three distance measures. The color and size of the circles are indicative of the number of species in the network (see Main). *r*_*i*_ = 0:4; *s*_*i*_ = 1:5; *c*_*i*_ = 1.

S2 The effect of multiple shifts on Euclidian distance *δ*, relative Shannon index change ∆s and the number of secondary extinctions. (a-b) *δ* and ∆*s* for real and random networks, as well as the ratios δ_real_/ δ_random_ and ∆*s*_real_= ∆*s*_random_. A positive ratio indicates a larger value in real networks and a negative ratio a larger value in random networks. (c) The number of secondary extinctions in real and random networks. *r*_*i*_ =0.4, *s*_*i*_ =1.5, *c*_*i*_ =1.

S3 The effect of multiple removals/extinctions on residual Euclidian distance *δ*^(−*k*)^, residual relative Shannon index change ∆s^(−*k*)^ and the number of secondary extinctions. (a-b) δ and ∆s for real and random networks, as well as the ratios δ_real_/δ_random_ and ∆s_real_/∆s_random_. A positive ratio indicates a larger value in real networks and a negative ratio a larger value in random networks. (c) The number of secondary extinctions in real and random networks. We did not observe any secondary extinctions with species removals, since there is facultative mutualism and species removals do not result in negative interactions. *r*_*i*_ =0.4, *s*_*i*_ =1.5, *c*_*i*_ =1.

S4 Relative Euclidian distance, relative Shannon index change and relative abundance change for different values of intra-specific competition. Left: weak competition (*s*_*i*_=1.1); Right: strong competition (*s*_*i*_ =3). *r*_*i*_ =0.4, *c*_*i*_ =1.

S5 Distance measures for single shifts in real-world vs. randomized networks for weak competition (*s*_*i*_ =1:.1). *r*_*i*_=0.4, *c*_*i*_ =1.

S6 Distance measures for single shifts in real-world vs. randomized networks for strong competition (*s*_*i*_ =3). *r*_*i*_ =0.4, *c*_*i*_ =1.

S7 Antagonism vs. extinctions for weak and strong intra-specific competition. Left: weak competition (*s*_*i*_ =1.1); Spearman’s *ρ* = −0.609 (p <10^−15^) for δ and *ρ* = −0: 475 (p <10^−15^) for ∆s. Right: strong competition (*s*_*i*_ =3);Spearman’s *ρ* = −0.563 (p <10^−15^) for δ. There is no significant decrease with degree centrality for ∆s, *ρ* = −0.0196 (*p* =0:242). *r*_*i*_ =0.4, *c*_*i*_ =1.

